# Phenologically Explicit Robustness Metric Reveals Increased Vulnerabilities in Temporal Plant-Pollinator Networks

**DOI:** 10.64898/2025.12.23.695469

**Authors:** Raoul Scribbans, Tim Rogers, Michael J.O. Pocock, Richard James

## Abstract

Robustness is a useful metric in evaluating the ability of an ecosystem to resist further extinction after species loss. This measure is straightforward to calculate from empirical interaction data, however doing so typically requires a static network of time-aggregated interactions, ignoring the finer-scale variation imposed by species phenologies. Although the analysis of time-aggregated ecological networks has been successful in capturing their overall structure, it often leads to the loss of information of the within-season variation that ecosystems typically exhibit. We demonstrate how this approach can also dramatically overestimate robustness by assuming all interactions are concurrent. Here, we develop a new measure of temporal robustness that incorporates the time-varying nature of species interactions. We apply our measure to plant-pollinator systems using both simulated and empirical networks, and obtain analytical predictions for the temporal robustness of randomly generated networks. In each case, our analysis reveals a substantial loss of information in the established static-network robustness measure, which obscures the crucial role of phenology in determining the vulnerability of pollinators to species loss. We find that pollinators active over long time periods appear more robust after time-aggregation, but typically have a much lower temporal robustness, indicating that long-lived pollinators may be more vulnerable to plant loss than previously thought.

## I. INTRODUCTION

Many ecosystems are under threat, owing to a wide range of environmental pressures. This is increasingly true for plant-pollinator systems [1], which are vulnerable to threats including loss of habitat (e.g. due to land use change) [2–4], increased use of pesticides [5, 6], and climatic changes such as drought or extreme weather due to global warming [7]. In order to safeguard these ecosystems and the pollination services they provide, an understanding of their ability to withstand species loss and maintain biodiversity is of great ecological importance. To this end, many studies have used a concept of ecosystem robustness (see e.g. [8–10]), quantifying an ecosystem’s ability to resist such pressures and avoid further extinctions after the loss of some of its species. An understanding of ecosystem robustness can therefore aid in the design of conservation strategies [11], and the implementation of effective ecosystem restoration [9, 12].

Although there are many ways through which one can quantify robustness [13], here we consider the node removal model introduced in [8] for plant-pollinator (bipartite, mutualistic) networks. In this model, species of one type are removed from the network, and species of the second type become extinct once they lose some (model-dependent) amount of their interactions. Here we limit considerations to the removal of plant nodes in a plant-pollinator network, but note that the analysis would be essentially identical for other bipartite mutualistic systems. Despite its widespread use as a network metric (see e.g. [9, 14]), a detailed analytical description of robustness did not appear until relatively recently in [15]. In this study, Jones et al. establish a number of useful results, showing that secondary extinction probability follows a hypergeometric distribution.

One element often overlooked in studies of network robustness is species phenology, and the variation of activity times within a season. It is well-established that phenology is a key determinant of network structure [16– 18]. By constraining interactions in time, phenology is a cause of forbidden links [19], and can explain much of the structure observed in real ecosystems [20]. It has been suggested that variation in phenology can increase species persistence by mediating competition [21], and that seasonal structure of plant-pollinator networks can act to increase their stability [22]. The shifting of phenologies, especially as a consequence of global warming, has been identified as causing phenological mismatch between species and loss of interactions [7, 23, 24]. Furthermore, several studies discuss how networks change over seasons and years, showing that virtually all network metrics, including connectance, nestedness, numbers of species, and species specialisation/generalisation are not fixed, but instead vary greatly throughout seasons [25–29].

Despite these considerations, studies which explicitly take phenology into account when evaluating network robustness (in the more general sense) are scarce. One example is given in [30], where Encias-Viso et al. consider a combination of within– and between-year dynamics to determine the long-term stability of mutualistic networks, and show that the distribution of species phenology has a large effect on the resilience of ecosystems to small perturbations. In [10], Ramos-Jiliberto et al. study an integro-difference equation model of plant-pollinator interactions throughout their lifecycles, and show that phenology is a strong determinant of species persistence. These exceptions aside, virtually all others consider a static interaction network, which is either a “snapshot” of the interactions over a short interval of time, or, more typically, a temporal aggregation of interactions over a longer sampling period. Time aggregation in this manner has been shown to misrepresent many key network metrics [26, 27], and in [31] Payrató-Borrás et al. show how in doing so we can lose non-trivial information about finer-scale network structure. These concerns have been raised previously, with multiple authors calling for a better framework through which to understand temporal ecological networks [32–34].

An illustrative example of the effects of time-aggregation on robustness is given in [26]. Here Caradonna et al. compare 12-15 weekly-aggregated networks with their season-long aggregation, and show how this aggregation misrepresents ecosystem structure, producing a network that tends to be more robust than its weekly components. This effect can be understood through Figure 1, where we show how this misrepresentation tends to inflate species degree and thus robustness. This idea is corroborated by [35], where a clear positive correlation is shown between pollinator (and plant) generality (the mean number of interactions per pollinator) and the temporal extent/scale of aggregation (from days to many years) of many empirical time-aggregated plant-pollinator networks. By incorrectly assuming that all interactions are concurrent, we obtain a network that is both more speciose and has more interactions at any given point in time, and thus appears more robust to species loss than one that is temporally resolved.

**FIG. 1:**
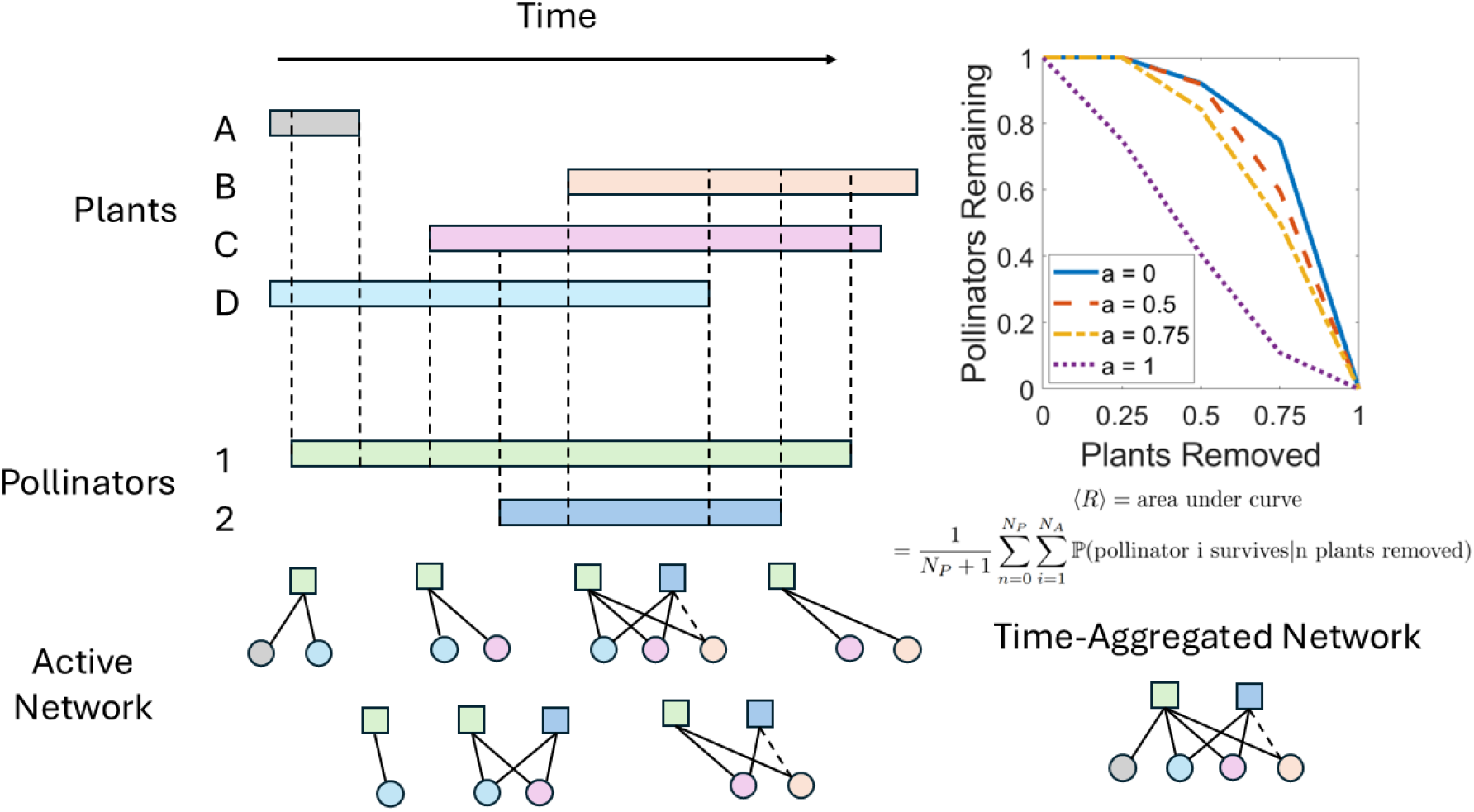
Illustration of example temporal network composed of *N*_*P*_ = 4 plants and *N*_*A*_ = 2 pollinators, with their respective active periods indicated in the top left. Each start/end of an active period delimits a pair of sub-intervals, each of which has a corresponding active network shown below. Here, solid lines indicate interactions, and dotted lines indicate co-occurrence without interaction. The time-aggregated network, shown in the bottom right, is the union of all the active networks and does not exist at any point in time. The extinction curves for this network, for various values of the MRT *a* (where the same value of *a* is assigned to both pollinators), are shown in the top right, with *a* = 0 corresponding to the time-aggregated network. The respective areas under them give an expected time-aggregated robustness of 11*/*15 ≈ 0.733 for *a* = 0, but a lower expected temporal robustness of 11*/*24 ≈ 0.458 for *a* = 1 (calculated using equation 1). Extinction curves are simulated with 1000 randomly chosen extinction sequences in each case. Adapted from [10].

Here, we introduce *temporal robustness*, a novel network metric which explicitly takes into account species phenology and the variation in interactions over time. In the following section we demonstrate how this can be calculated for a given plant-pollinator network, and how it differs from time-aggregated robustness. Furthermore, we show how (temporal) robustness can be disaggregated in terms of the *individual robustness* of each pollinator, which provides insight the effects of species loss on each pollinator in a given ecosystem. We then apply these metrics to empirical plant-pollinator networks, and demonstrate that time-aggregation acts to overestimate species robustness in real plant-pollinator systems. This effect is especially strong for pollinators active over long times, suggesting that these pollinators may be much more vulnerable to plant loss than predicted by time-aggregated models. In section III we develop the *uniform model*, a simple and illustrative temporal network model. Using results from the theory of random interval graphs, we obtain analytical expressions for both the temporal and time-aggregated robustnesses of these model networks, providing insight into the general features that make temporal networks robust. In section IV we then compare the uniform model with empirical networks, and show that real plant-pollinators systems appear to be much more temporally robust than expected from simulations. We also introduce the *linear interaction model*, in which long-lived pollinators are much more generalist than those that are short-lived. This model captures observed pollinator degree distributions well, suggesting that, by being more generalist, long-lived pollinators are able to maintain a relatively high temporal robustness.

## II. TEMPORAL ROBUSTNESS

We begin with the robustness model first considered in [8]. Here, plant nodes are sequentially removed from a plant-pollinator network via *primary extinctions*. In a time-aggregated (static) network, we say that a pollinator will undergo a *secondary extinction* once all of its interactions have been lost. By plotting the proportion of pollinators remaining against the proportion of plants removed, one obtains the *extinction curve* for a given *extinction sequence* of plants, illustrated in Figure 1 (top right). The area under this curve determines the network’s robustness to plant loss for a given sequence. By averaging over randomly chosen extinction sequences one obtains the expected robustness. In what follows we always average over randomly chosen extinction sequences, and therefore only consider expected robustness.

A useful way of expressing robustness is as the average *individual robustness r* of each pollinator. The individual robustness of a given pollinator is equivalent to the expected proportion of plants that must be removed for it to go extinct. This therefore describes the effects of node removal on an individual pollinator (rather than the ecosystem as a whole), providing information about its vulnerability to plant loss.

In order to define the temporal robustness of a temporally resolved network, we first determine the *active period* of each species. This corresponds to the period over which a given species is able to interact, and represents either a blooming period (for plants) or a flight period (for pollinators). Interactions can then only occur between a plant and pollinator during the intersection of their respective active periods, and will therefore be forbidden if the active periods do not overlap. The *active network* at a given point in time can then be found by taking the set of plants, pollinators and interactions which are active at that time. The corresponding time-aggregated network instead has all plants, pollinators and interactions active, and is the union of all the active networks. This is illustrated in Figure 1 for a model temporal network composed of two pollinators and four plants.

Given a pollinator’s active period, it is then necessary to define the conditions under which it undergoes a secondary extinction. Here we assume that a pollinator with an active period of duration *w*_*A*_ requires interactions with plants (i.e. food) for some continuous proportion *a* ∈ [0, 1] of that active period in order for it to survive. The duration *aw*_*A*_ can then represent the amount of time it takes for an individual of the species to successfully reproduce (e.g. in the case of solitary bees), or for a colony to provision enough food for the season (e.g. for social bees), and for the species as a whole to survive to the next generation. We therefore refer to the proportion *a* as the *minimum reproduction time* (MRT) for a given species of pollinator.

For *a* = 1 we therefore have that a pollinator requires interaction over its entire active period, and will become extinct if there are any temporal gaps in the plants it interacts with. If instead *a* ≪ 1, it becomes likely that any of the pollinator’s interactions is sufficient for it to survive. In this case we have that all interactions must be removed for the pollinator to go extinct, and thus return to the time-aggregated case. We therefore say that having *a* = 0 for all pollinators is equivalent to the time-aggregated case. Intermediate values of *a* will then interpolate between these two extremes.

For a given value of *a* we can then determine the *extinction conditions* for each pollinator. These are the sets of plants that, if lost, will cause a given pollinator to go extinct. For example, if pollinator 1 of the network illustrated in Figure 1 has an MRT *a* = 1, it will go extinct if plant *D* is removed, or if both plants *B* and *C* are removed. Given these conditions we can calculate analytically the expected individual temporal robustness for a given pollinator as

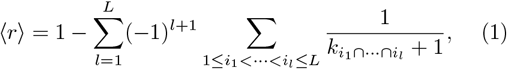

where *L* is the number of extinction conditions the pollinator has, and 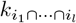 is the number of unique plants that must go extinct for extinction conditions *i*_1_, …, *i*_*k*_ to all be fulfilled. This can be averaged over pollinators to obtain the temporal robustness of the whole network. Details of this calculation and the determination of extinction conditions can be found in section S1 of the Supplementary Material. This approach allows one to calculate the robustness for any set of non-independent extinction conditions (i.e., those based on phenology or otherwise).

For a time-aggregated network (*a* = 0), a given pollinator of degree *k* has a single extinction condition, namely that all plants it interacts with are removed. In this case equation 1 reduces to ⟨*r*⟩ = *k/*(1 + *k*), and we recover the result for time-aggregated robustness,

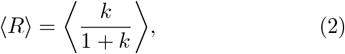

given in [15].

Alternatively, one can simulate plant extinctions to obtain an extinction curve for a given network. A given pollinator will remain after a set of plants has been removed if none of its extinction conditions have been fulfilled. By simulating over many (randomly chosen) extinction sequences of plants we can obtain an extinction curve (see Figure 1), and numerically evaluate the area under it to obtain the robustness of the network. This can be preferable if the number of extinction conditions becomes very large, and equation 1 becomes slow to evaluate.

Given a set of plants and pollinators, their respective active periods, and their interactions (e.g., as a static adjacency matrix), we can therefore calculate the (temporal) robustness of the corresponding temporal network. Here we have done so for 8 yearly networks based on data gathered in open pine forest ecosystems in Doñana Natural Space, southern Spain, from 2015 to 2022, provided in the study [36]. In Figure 2 we illustrate an example yearly temporal network (data gathered in 2022), and show the active periods (A) and interactions (B) for that year. Using this information to define a temporal network, we can evaluate the corresponding robustness via equation 1.

**FIG. 2:**
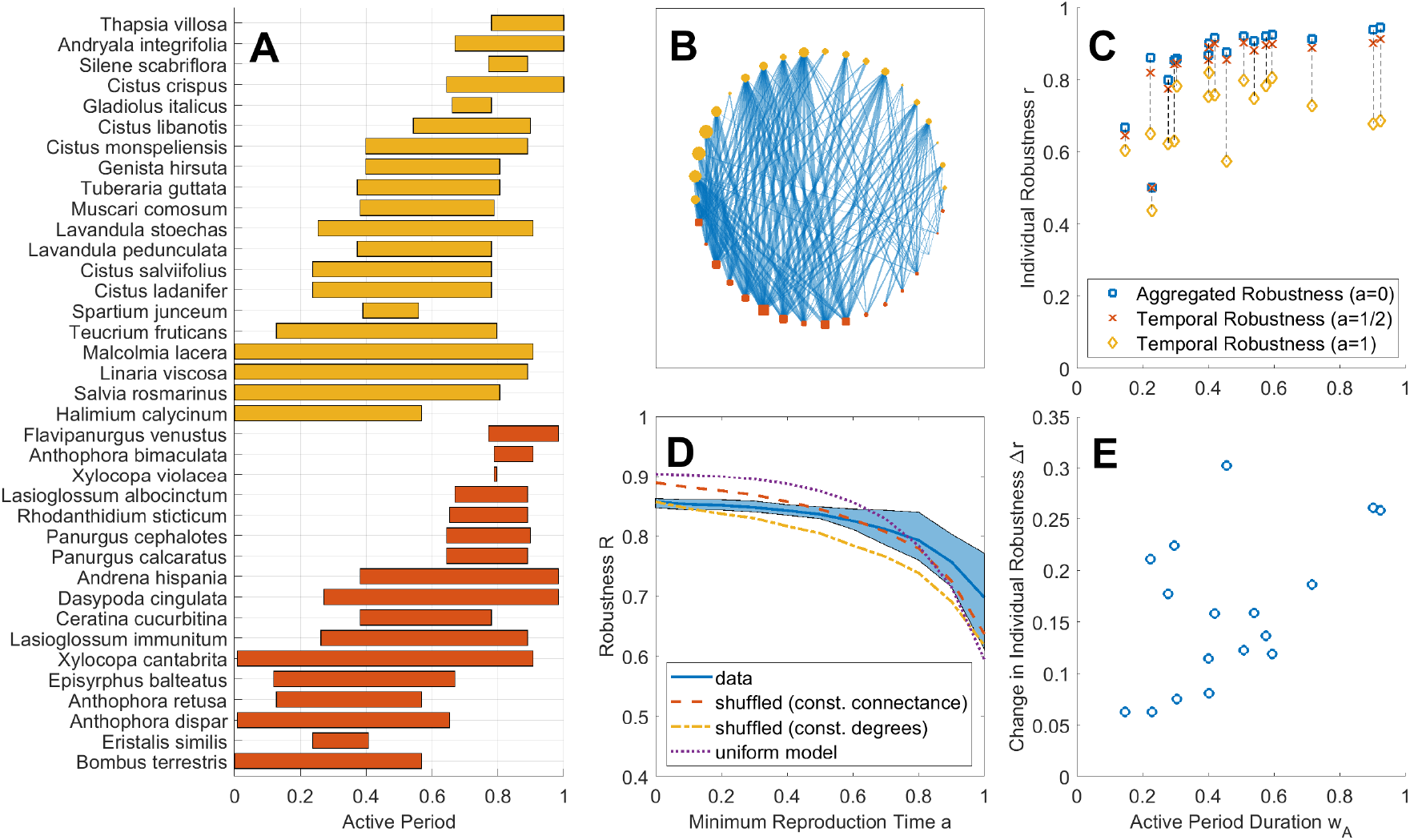
Example phenologies (A) and network (B) corresponding to a single year (2022) from the Doñana dataset. In panel A active period durations represent the proportion of the whole flowering season (118 days total) that the given species was active. Panel B shows (time-aggregated) interactions, where species are ordered by midpoint of active period for pollinators (red, lower) and plants (yellow, upper). Here node size is proportional to interval duration and link width is proportional to interaction (active period overlap) duration. In panels C, D, and E values are instead averaged over all 8 years. Panel D shows robustness against MRT *a* (blue line), with the shaded blue region representing the minimum and maximum yearly network robustnesses for each value of *a*. These are compared to two different randomisations of the networks, which randomly reassign interactions whilst observing phenological constraints and either maintaining total number of interactions (red) or the degree of each pollinator (yellow). They are also compared to the uniform interval model (introduced in section III), with parameters inferred from data. Whilst the real networks are not as robust (on average) as the others for low *a* (especially for time-aggregated robustness at *a* = 0), they tend to be the most temporally robust for high *a*. In panel C we show the individual robustness of the pollinators. Individual robustness is calculated for *a* = 1 (yellow diamond), *a* = 0.5 (red cross) and *a* = 0 (blue square). Panel E indicates the corresponding change in robustness due to time-aggregation, i.e. Δ*r* = *r*(*a* = 0) − *r*(*a* = 1), which correlates positively with pollinator active period duration.

We note here that we have repeated this analysis (and much of what follows) for 6 other temporal plant-pollinator networks, which can be found in section S3 of the Supplementary Material. In the main text we focus on the Doñana dataset however, as phenologies were given explicitly, and we are able to average results over the 8 years. Results are, in general, qualitatively similar between datasets, and those illustrated here therefore offer a representative summary of our findings.

From these networks we can observe how the introduction of phenology has a substantial effect on ecosystem robustness. In panel D of Figure 2 we show that in each year of the Doñana dataset there is a reduction in ⟨*R*⟩ as the MRT increases from *a* = 0 to *a* = 1 (where all pollinators have been assigned the same value of *a*). We also demonstrate how the real networks compare to various randomisations of the data. We see that, whilst the empirical networks are not particularly robust for lower values of *a*, they are (on average) more robust than the randomisations for larger values (*a* ⪆ 0.8). This suggests that real plant-pollinator networks are arranged to have a high temporal (but not necessarily time-aggregated) robustness, further indicating that time-aggregated robustness may be a less important metric than temporal robustness for ecological networks which vary over time.

We can also use equation 1 to evaluate the individual robustness of each of the pollinators in a given network. This is shown in panel C of Figure 2, where we have calculated the individual robustness of each pollinator for various values of the MRT *a*. Here, the time-aggregated robustness tends to increase with pollinator active period duration *w*_*A*_. This is due to (time-aggregated) degree: pollinators which are active for longer periods tend to have more interactions, and thus a higher time-aggregated robustness. In the temporal (*a* = 1) case however, the relationship is less clear. We can however observe a positive correlation between pollinator active period duration and the change in robustness, i.e. the difference between time-aggregated (*a* = 0) and temporal (*a* = 1) robustnesses, which is shown in panel E of Figure 2. This suggests that the overestimation in robustness as a consequence of time-aggregation is more pronounced for longer-lived pollinators. This result is intuitive, as pollinators active over longer times have more scope for variation in interactions, and also more opportunities to go extinct due to species and/or interaction loss. This highlights a potential vulnerability of longer-lived species, which a time-aggregated model is likely to overlook.

Whilst the methods illustrated here are useful for evaluating the robustness of a given temporal network, or of the individual robustnesses of their constituent species, they do not provide much information about the general features which make a network robust. In the following section we therefore introduce a simple random network model, for which we can estimate temporal robustness based on a small number of parameters.

## III. ROBUSTNESS OF RANDOM TEMPORAL NETWORKS

Here we introduce the *uniform model*: a simple model of temporal plant-pollinator networks. We construct a network instance by taking *N*_*A*_ pollinators, and assigning each of them an interval of duration *w*_*A*_ and MRT *a*. Similarly, we take *N*_*P*_ plants and assign each with an interval of duration *w*_*P*_. Each interval is then placed randomly within [0, 1], which represents the full flowering season, such that the start of each pollinator (plant) interval is drawn uniformly from [0, 1 − *w*_*A*_] ([0, 1 − *w*_*P*_]). Each interval represents the proportion of the season for which a given species is active. Any plant-pollinator pair with overlapping intervals will then interact with a probability *p*_*int*_ ∈ [0, 1], which determines the expected connectance of the active network at a given point in time. An example uniform model network is illustrated in panels A and B of Figure 3.

**FIG. 3:**
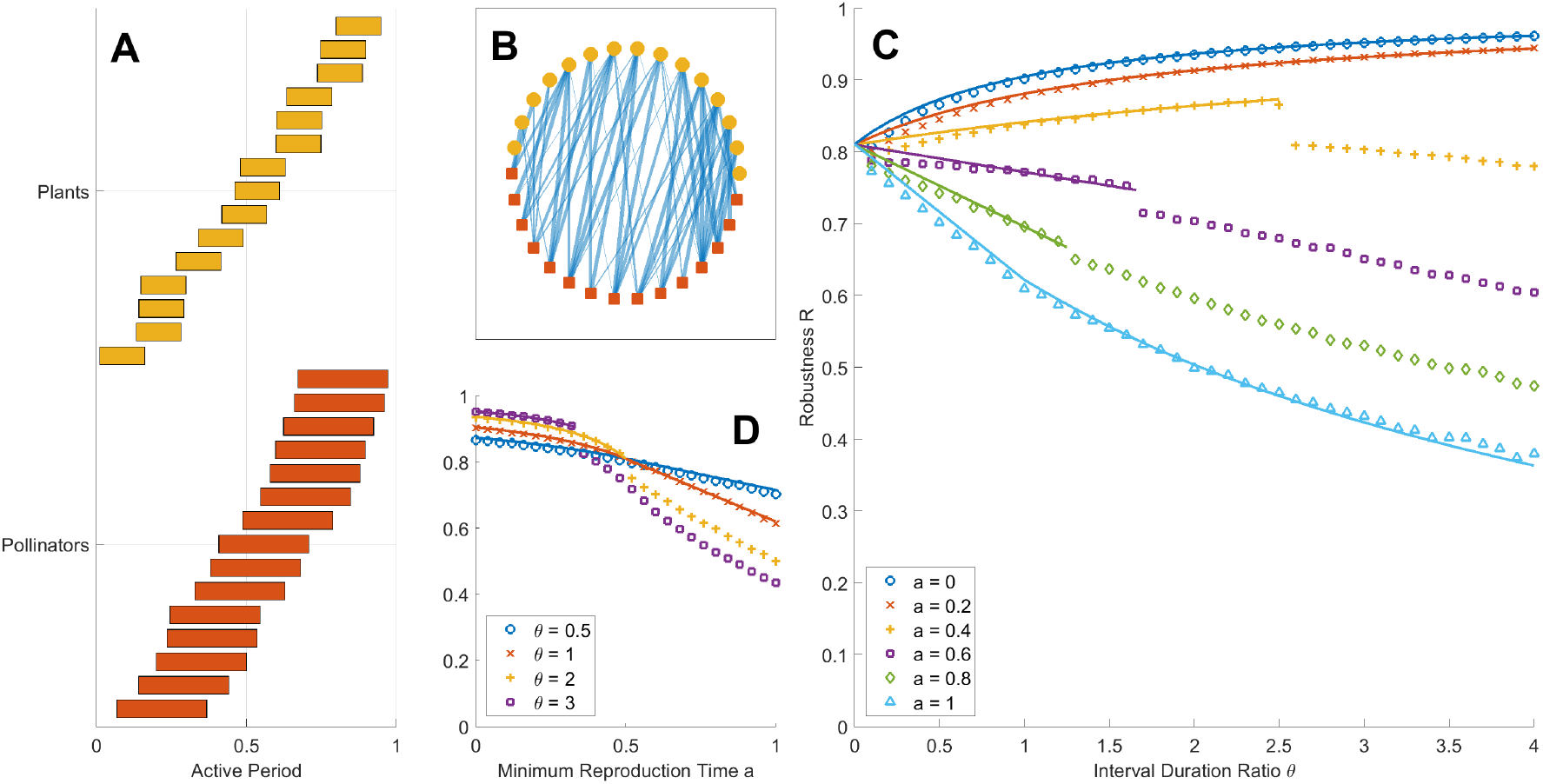
A temporal network, randomly generated as described in section III, is illustrated panels A and B. Panel A illustrates the active periods of both plants (top, yellow) and pollinators (bottom, red), with species ordered by active period time. Species interactions are shown the panel B, where nodes are also ordered from left to right by active period time for both pollinators (red squares) and plants (yellow circles), and link width is proportional to interaction (active period overlap) duration. A comparison between simulation and prediction of the temporal robustness of random networks is shown in panels C and D. Each point corresponds to the average robustness of 100 network instances, each evaluated numerically over 100 extinction sequences. Solid lines correspond to approximations from equations 3 (*a* = 1), S27 (*a* = 0), S28 (1*/*2 ≤ *a* ≤ 1, *aθ <* 1), or S29 (0 ≤ *a* ≤ 1*/*2, *aθ <* 1). Discontinuities in the robustness can been seen at *aθ* = 1 (see main text for details). In both cases plant interval duration is fixed at *w*_*P*_ = 0.05. Other parameters used are *N*_*P*_ = *N*_*A*_ = 200, and *p*_*int*_ = 0.5, giving *ρ* ≈ 5.263. For visual clarity, the network shown in the panels A and B was instead generated with *N*_*P*_ = *N*_*A*_ = 15, *w*_*P*_ = 0.15, *w*_*A*_ = 0.3, and *p*_*int*_ = 0.5.

For the case *a* = 1, a pollinator can only survive if the plants it interacts with fully cover its interval. Using results from random interval graph theory [37], we obtain approximate expressions for this covering probability, allowing us to calculate the expected temporal robustness for a random network instance. Details of these calculations, and others in this section, can be found in section S2 of the Supplementary Material. The result is a polynomial in the ratio of the interval durations *θ* = *w*_*A*_/*w*_*P*_

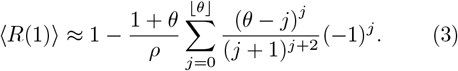

Here we write *R* = *R*(*a*) as a function of the MRT *a*, and note that we are now averaging over network instances in addition to extinction sequences. We also introduce the effective plant density *ρ* = *p*_*int*_*Nw*_*P*_ */*(1 − *w*_*P*_), which determines the expected number of plants that will interact with a pollinator at a given point in time. We assume here that *θ*, and therefore the number of terms in the equation above, is of order 1.

The pollinators in this model have an approximately binomial degree distribution. Using this and equation 2 we then obtain an expression for the expected time-aggregated robustness (*a* = 0), given by equation S27 of the Supplementary Material. We also obtain estimates for intermediate values of *a* in the case *aw*_*A*_ < *w*_*P*_, given by equations S28 (for *a* ≥ 1*/*2) and S29 (for *a* ≤ 1*/*2).

Comparison of these approximations to simulation are illustrated in panels C and D of Figure 3, with good agreement between theory and simulation. Since all pollinators are drawn identically, the results here describe both the expected robustness of an ecosystem of *N*_*A*_ pollinators, and the expected individual robustness of a single pollinator. Both plots show how expected robustness reduces monotonically with the MRT *a*. We also see that this effect is exacerbated for larger values of *θ*, i.e. for pollinators with longer active periods. In panel C of Figure 3 we observe that, as the pollinators’ active periods increase, the time-aggregated robustness increases, but the temporal robustness decreases. Similarly to the empirical networks shown in Figure 2 in the previous section, a pollinator active over a long time is likely to have a higher degree, and thus a higher (individual) time-aggregated robustness (according to equation 2). However, it is also likely to have more routes to extinction, reducing its temporal robustness. This helps to explain the correlation between the change in robustness (from temporal to time-aggregated) and pollinator active period duration described by panel E of figure 2, and suggests that pollinators active over long periods are likely to be much more vulnerable to species loss than predicted by time-aggregated models.

Discontinuities are shown in panels C and D of Figure 3, which occur at *aθ* = 1. These correspond to the threshold at which a single plant interval is sufficient (*aθ* ≤ 1), or not (*aθ >* 1), to cover enough of a pollinator’s interval for it to survive. This effect is a consequence of having all plants (pollinators) with the same interval duration *w*_*P*_ (*w*_*A*_), and the introduction of a distribution in either (or both) plant or pollinator interval duration would eliminate these discontinuities.

In [15], Jones et al. show that, for a given number of interactions, time-aggregated robustness is maximised when all pollinators have (as close as possible to) the mean degree. This can be understood through Jensen’s inequality – since *k/*(1 + *k*) is a concave function of *k* we have that

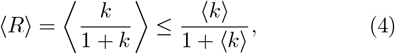

with equality occurring where *k* = ⟨*k*⟩ . Similarly, equation S27 is a concave function of *θ*. This means that, for a given plant interval duration *w*_*P*_ and mean pollinator interval duration ⟨*w*_*A*_ ⟩, expected time-aggregated robustness is maximised when all pollinators are of equal interval duration *w*_*A*_ = ⟨*w*_*A*_ ⟩. In contrast, equation 3 describes temporal robustness as a convex function of *θ* (for *θ >* 1). This means that a distribution in pollinator active period duration will act to increase temporal robustness, with ⟨*R*(1) ⟩ instead being minimised when all pollinators are of equal interval duration. This may help to explain the wide variation in degree and active period duration typically observed in real ecosystems [20].

In the following section we return to the temporal networks inferred from the Doñana dataset, and compare them to the uniform model developed here. We show that, whilst the uniform model is successful in capturing some features of the real ecosystems considered, it also differs in meaningful ways. We therefore demonstrate how this model can be refined and extended.

## IV. COMPARISON OF EMPIRICAL AND MODEL TEMPORAL NETWORKS

The uniform model introduced in the previous section suggests that the individual robustness of pollinators active over longer periods ought to be higher for time-aggregated networks, but lower for temporal networks (when *a* is high). We have shown in Figure 2 that the former appears true for the data considered here, as pollinators with larger interval durations tend to have a higher time-aggregated robustness. In Figure 4 we reproduce this result with a comparison to the uniform model (yellow lines) introduced in the previous section, for both time-aggregated (*a* = 0, left) and temporal (*a* = 1, right) robustnesses, where uniform model parameters have been estimated from the data. We observe that, in the uniform model, time-aggregated robustness tends to increase similarly to that of the real data. This is driven by pollinator degree: in the lower panel we show that, in both the data and the uniform model, pollinators active over longer periods tend to have more interactions, and therefore have an increased individual (time-aggregated) robustness *r* = *k/*(1 + *k*).

**FIG. 4:**
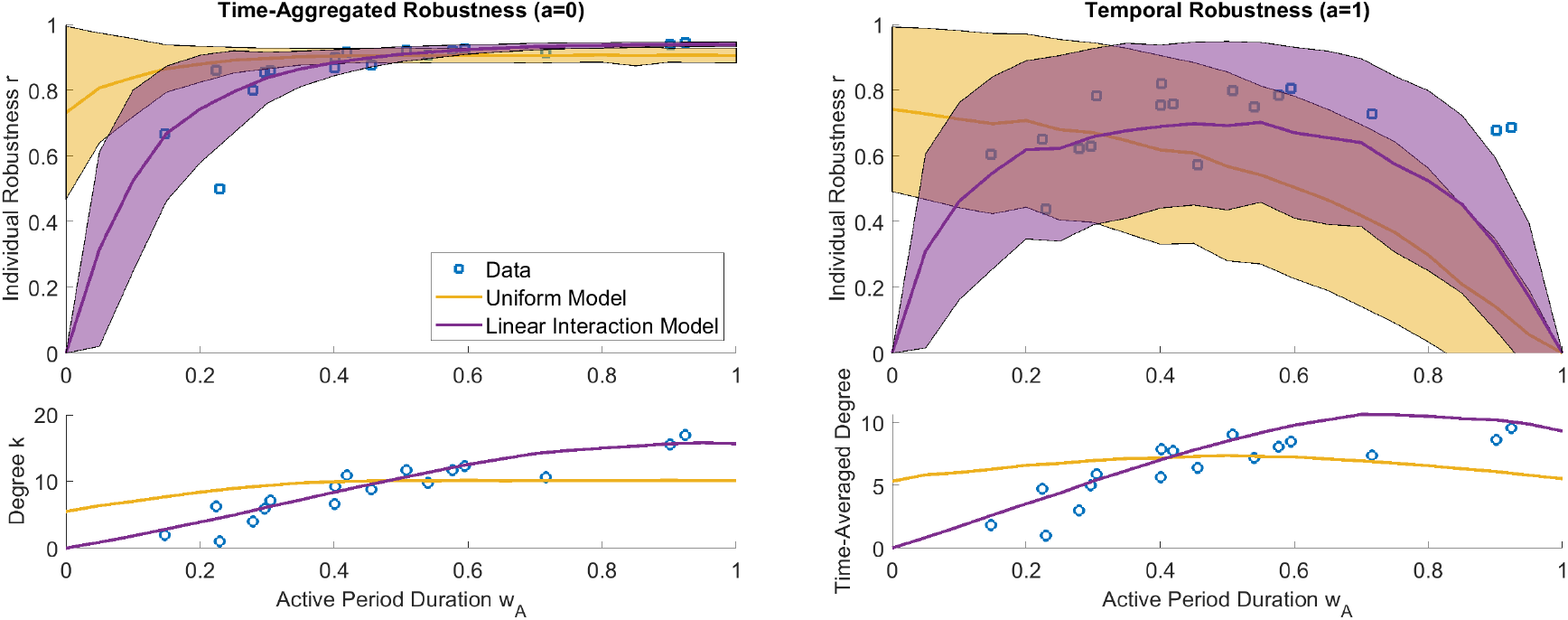
Comparison of Doñana dataset to the uniform model introduced in section III (yellow) and the linear interaction model introduced in section IV (purple). Upper panels show individual robustness, with solid lines showing mean values, and shaded regions indicating standard deviation, of 100 randomly generated network instances for each pollinator interval duration *w*_*A*_ and MRT *a*. The lower panels compare the mean degree (left) and mean time-averaged degree (right) of each pollinator in the generated networks to those of the data. Parameters used are *N* = 20, *M* = 17, and *w*_*P*_ = 0.5504 (uniform model). In the linear interaction model the constant of proportionality between interaction probability and overlap duration is 1.5093, and plant widths are drawn from a beta distribution with parameters *α* = 3.5463 and *β* = 2.9000.

The uniform model also predicts that temporal robustness will decrease monotonically with interval duration *w*_*A*_, dropping off significantly as *w*_*A*_ approaches 1, however neither of these predictions is reflected in the data. This is unsurprising, indicating that real ecosystems, and particularly the interactions of longer-interval pollinators, tend to be arranged so as to be much more temporally robust than expected from the null (uniform) model.

One way pollinators active over long times are able to remain robust is illustrated by the lower-right panel of Figure 4, where we show that (in addition to total degree) time-averaged degree (the mean number of interactions a pollinator has over its active period) is correlated with interval duration. This suggests that longer-lived pollinators are inherently more generalist than shorter-lived ones, rather than simply having more interactions because they are around for longer. Through this generalism they are therefore able to avoid the drop-off in robustness exhibited by the uniform model.

In order to better account for this we introduce the *linear interaction model*. In this model, the probability a given plant-pollinator pair interact is directly proportional to the duration of their overlap. This relationship is also reflected in the data, and we are therefore able to estimate an empirical constant of proportionality between overlap duration and interaction probability. We also infer a probability distribution of plant interval durations by fitting a beta distribution to those in the data. Although it has been suggested that a plant interval durations tend to follow a log-normal distribution [20], we instead fit a beta distribution here, which describes observed distributions well, with the added advantage of constraining active period durations within [0, 1]. Any plants in the linear interaction model will then have an interval duration *w*_*P*_ drawn randomly from this beta distribution. Further details of the inference of both plant interval distributions and interaction probabilities are given in section S4 of the Supplementary Material.

Through the linear interaction model we obtain the purple lines and regions in Figure 4. This offers a closer representation of the data than the uniform model, and better captures the relationships between total degree/time-averaged degree and interval duration. It also predicts that temporal robustness will be maximised for an intermediate value of interval duration, rather than for *w*_*A*_ → 0. Here, pollinators active over very long times are less robust due to the increased probability of a gap in their interactions, whereas those active over very short times are less robust due to their lower number of total interactions. This is also shown in the data, however the longest-interval species are significantly more temporally robust than the model predicts (largely due to the hard boundaries imposed at the start and end of the season in either model). Interestingly, the introduction of a plant width distribution has virtually no effect on the uniform model, however in conjunction with the linear interaction probabilities it produces an increase in robustness for the longer-interval pollinators who are able to take advantage of longer-interval plants. This also helps to explain why longer-interval pollinators remain relatively temporally robust.

## V. DISCUSSION

We have introduced temporal robustness as a refinement of the time-aggregated robustness widely used in studies of plant-pollinator networks. A consistent theme here is that the explicit inclusion of species phenology can, and often does, have a large effect on the measure of robustness of plant-pollinator networks to species loss. This is true of both model networks, and those based on real data. This suggests that studies which aim to measure species interactions ought to do so over sufficient time, and sufficiently often, to be able to capture the full temporal structure, and therefore robustness, of a given ecosystem.

A further useful concept introduced here is individual robustness. This metric offers a way to identify particularly vulnerable species, which could aid in conservation efforts (especially by considering those pollinators’ extinction conditions). Through this, we have illustrated that the decrease in robustness introduced by phenology is likely to be more severe for longer-interval pollinators. That said, pollinators in real ecosystems are able to maintain a relatively high individual robustness by being more generalist and having long lasting interactions. Real ecosystems appear much more temporally robust than would be expected from the two null models introduced here, and it therefore appears likely that temporal robustness is a more useful concept than its more commonly used time-aggregated counterpart.

Sampling issues are likely to over-represent certain species and interactions however, especially those species which have longer active periods. Furthermore, species which coexist for longer are more likely to be observed interacting than those who do so briefly, which may explain the observed correlation between interaction probability and overlap duration. In reality, it seems likely that there is a combination of sampling effects and co-ordination of species phenologies producing the networks observed here. Therefore, it may be true that longer-lived pollinators are more vulnerable than our data analysis suggests, with a drop-off in robustness similar to that of the uniform model occurring. That said, shorter-lived interactions are likely to have less of an effect on temporal robustness. In fact, the study conducted to obtain the Doñana dataset focussed on species that were present over all 8 years of the study. Therefore, it is likely that longer-lived species which are easier to detect are further overrepresented here. The networks discussed in section S3 have more pollinators with short active periods, which also tend to have a smaller difference between aggregated and temporal robustness.

The model of temporal robustness introduced here clearly simplifies a lot of ecological detail. Many studies of ecosystem robustness have considered extensions to the basic node removal model, including varying interaction strength [15, 38], non-random extinction orders [8, 39], and extinction cascades [38], where secondary pollinator extinctions lead to further plant extinctions and so on. Whilst these effects have been understood with regard to time-aggregated networks, it would be interesting to study the interplay between them and temporal structure, and whether these extensions exacerbate or ameliorate the loss in robustness due to time-structure we have observed here. It would also be interesting to see if extinction cascades are localised in time, or if they propagate through the whole temporal network. Furthermore, the temporal framework developed here could be extended to more complex phenologies, such as those of multivoltine species, or to study the temporal variation in species abundance.

The random network models introduced here are also relatively simple, and, whilst successful in capturing many features of the real data (see e.g. Figure 4), could be further refined. For instance, we have only considered a uniform distribution of plant and pollinator interval start times. In reality we expect some variation in this, which would likely lead to more (or less) dense periods of interaction, which are likely to be more (or less, e.g. the so-called “June gap” observed by British beekeepers) robust [40]. Furthermore, in a real ecosystem we may expect to observe some coordination between plant and pollinator active periods. This is illustrated here by the solitary bee *Flavipanurgus venustus*, which is known to be monolectic on *Cistus crispus* [41], and is the least robust pollinator (for all values of *a*) shown in Figure 4. Consequently, its maximal degree of *k* = 1 leads to maximal individual robustness of *r* = 1*/*2 (for all *a*). In order for *Flavipanurgus venustus* to survive, the active periods of the two species must therefore be well-aligned, which, indeed, they are in the data. The linear interaction model accounts for this to an extent, by ensuring that interactions between long-overlapping species are more likely, however it does not affect the probability that a given plant-pollinator pair will overlap. This may explain why the pollinators in the Doñana dataset are, on average, more temporally robust than those of model the predictions.

Whereas shorter-lived pollinators may be able to coordinate their periods of activity with the plants they interact with, especially those pollinators which are oligolectic (or monolectic), this is difficult for pollinators active over longer time periods. For example, bumblebees, such as *Bombus terrestris*, can survive through their sociality.

Despite the long active period of the colony as a whole (≈ 0.9 yearly average in the Doñana dataset), an individual will be expected to live for a much shorter time. Furthermore, the requirement that a continuous proportion of the active period is covered is less stringent for social pollinators, as the survival of the colony does not depend as strongly on the survival of a given individual. Therefore, we can expect social pollinators such as *Bombus terrestris* to be more accurately modelled by a relatively low value of the MRT *a*. Indeed, many pollinators will adopt strategies through which they can avoid extinction despite a lack of floral resources, such as aestivation, or by producing multiple broods within a given season, thereby effectively reducing their MRT.

The simplicity of the models introduced here is advantageous however, in that they are easily applicable to data, allowing us to capture information about empirical ecosystem structure. That said, there are relatively few empirical studies that include both species interactions and phenologies. In light of the large effects we have shown phenology to have on ecological network structure and robustness, it seems pertinent that more work is done in this area, as suggested by others previously [32–34]. Despite the relatively consistent results we have seen across networks, it would be interesting to see if the phenological patterns we have observed are also seen across other ecosystems.

## ACKNOWLEDGMENTS

This work was supported by funding from Outlook Energy. We would also like to thank Nerea Montes-Perez for providing early access to the Doñana dataset used here.

