## Supplementary Material for "Phenologically Explicit Robustness Metric Reveals Increased Vulnerabilities in Temporal Plant-Pollinator Networks"

Raoul Scribbans[1], Tim Rogers[2], Michael J.O. Pocock[3], Richard James[1]
[1] *Department of Physics, University of Bath, Bath, BA27AY, UK.*
[2] *Department of Mathematical Sciences, University of Bath, Bath, BA27AY, UK. and*
[3] *UK Centre for Ecology & Hydrology, Wallingford, Oxfordshire, UK*

### S1. CALCULATING EXPECTED ROBUSTNESS FROM EXTINCTION CONDITIONS

For the case that $a = 1$ the extinction conditions of a particular pollinator of degree $k$ can be found as follows: First, each start/end of a plant interval that falls inside a pollinator's active period will divide it into two separate regions. Let the number of these regions be $L$, and the number of interacting plants overlapping with each region be $k_l$, where $l = 1, \ldots, L$ labels the region and $0 \leq k_l \leq k$. Each region is then assigned a corresponding "extinction condition" $E_l$, which is the event that the $k_l$ interacting plants are removed (or go extinct). The event that the pollinator is extinct is then equivalent to the event that at least one of the conditions $\{E_l\}$ is fulfilled. The probability that the condition $E_l$ has been fulfilled at step $j$ (i.e. once $N - j$ plants have been removed) is given by $\binom{j}{k}/\binom{N}{k}$. Since a given extinction condition may be a subset/superset of others, it can be more useful to consider the set of "minimal" extinction conditions. This is the set of extinction conditions $\{E^{min}\} = \{E_l | E_l \not\subset E_m \forall m \neq l\}$, i.e. those that are not subsets of any other conditions (or, equivalently, the set of interacting plants in the corresponding region of the plant interval is not a superset of the set of interacting plants in another region of the interval). In the case that any section of the pollinator interval does not coincide with any interacting plant intervals, the pollinator has no extinction conditions and is instead "unconditionally extinct".

if $0 < a < 1$, we can instead consider a window of width $aw_A$ placed such that its left-most edge is at the start of the pollinator interval. The pollinator will survive if the entirety of this window is covered by interacting plants and the extinction conditions for this window can be found as before. We can then slide this window along until either of its edges reach the start or end of a plant interval, and the extinction conditions for the window change, until the right edge of the window reaches the end of the pollinator interval. For a pollinator to go extinct, we must have that the extinction conditions of all window positions are fulfilled, and so the total set of extinction conditions is given by the union of the extinction conditions of each window position. The minimal extinction conditions can then be found from these conditions as before.

if $a = 0$, any interaction is sufficient for the pollinator to survive, and we recover the time-aggregated case. In this case the single extinction condition is the event that all of its interacting plants have been removed.

Given a set of minimal extinction conditions, we can calculate the expected robustness, averaged over extinction sequences, starting from the definition

$$\langle R \rangle_S = \frac{1}{M(N+1)} \sum_{i=0}^{N} \langle m_i \rangle_S, \tag{S1}$$

where $\langle m_i \rangle$ is the expected number of pollinators remaining after step $i$. This can be written

$$\langle m_i \rangle_S = M - \sum_{j}^{M} p_{ex,j,i}, \tag{S2}$$

where $p_{ex,j,i}$ is the probability that pollinator $j$ has gone extinct after step $i$. $p_{ex,j,i}$ can be calculated as the probability of the union of the pollinator's minimal extinction conditions:

$$p_{ex,j,i} = \Pr\big(\bigcup_{l=1}^{L_j} E^{(j)}_{min,l} | i\big), \tag{S3}$$

which can be rewritten according to the inclusion-exclusion principle as

$$p_{ex,j,i} = \sum_{l=1}^{L_j} (-1)^{l+1} \sum_{1 \leq i_1 < i_2 < \cdots < i_l \leq L_j} \Pr\big(E^{(j)}_{min,i_1} \cap E^{(j)}_{min,i_2} \cap \cdots \cap E^{(j)}_{min,i_l} | i\big), \tag{S4}$$

where $L_j$ is the number of minimal extinction conditions pollinator $j$ has. In the expression above, for each value of $l$, the second summation is over all combinations of $l$ extinctions conditions. The probability that an intersection of extinction conditions occurs at step $i$ is given by the combinatorial factor

$$\Pr\left(E^{(j)}_{min,i_1} \cap E^{(j)}_{min,i_2} \cap \cdots \cap E^{(j)}_{min,i_l} | i\right) = \binom{i}{k_{i_1\cap\cdots\cap i_l}} \Big/ \binom{N}{k_{i_1\cap\cdots\cap i_l}}, \tag{S5}$$

where $k_{i_1\cap\cdots\cap i_l}$ is the number of unique plants that are required to go extinct for conditions $i_1, ..., i_l$ to be fulfilled. Substituting this into equation S1, and using that $\sum_{i=0}^{N}\binom{i}{k} = \binom{N+1}{k+1}$ then gives

$$\langle R\rangle_S = 1 - \frac{1}{M}\sum_{j=1}^{M}\sum_{l=1}^{L_j}(-1)^{l+1}\sum_{1\le i_1<i_2<\cdots<i_l\le L_j}\frac{1}{k_{i_1\cap\cdots\cap i_l}+1}. \tag{S6}$$

The above equation is valid for any given instantiation of a temporal network, and any plant-pollinator network in which the pollinators have an arbitrary number of extinction conditions.

### S2. ESTIMATING ROBUSTNESS OF A MODEL ECOSYSTEM

Here we derive estimations for the robustness (both temporal and time-aggregated) of the random network model introduced in section S2 of the main text. The results here are valid for both the robustness of a network composed of $M$ identical pollinators, or the individual robustness of a single pollinator.

For interval widths $w_A + 2w_P \le 1$, pollinators can be said to be either in the "bulk" of the interval (with midpoint $t \in [w_P + w_A/2, 1 - w_P - w_A/2)$ or at the edges ($t \in [w_A/2, w_P + w_A/2]$ or $t \in [1 - w_P - w_A/2, 1 - w_A/2]$). For a given pollinator with interval midpoint $t$, there is a constant probability that a randomly chosen plant will interact with it, and so its degree distribution will be binomial. Those in the bulk will interact with a randomly chosen plant with probability $p = p_{int}(w_A + w_P)/(1 - w_P)$, and those at the leftmost edge will do so with probability $\tilde{p}(t_s) = p_{int}(t_s + w_A)/(1 - w_P)$, where $t_s$ is the pollinator start time. Those at the rightmost edge will have a symmetrical degree distribution to those at the left, and the pollinator degree distribution can therefore be written

$$\begin{aligned} p_A(k) = (1 - (w_A + 2w_P))P_{bin}(k, N, p) \\ +2\int_0^{w_P} P_{bin}\left(k, N, \tilde{p}(t_s)\right) dt_s, \end{aligned} \tag{S7}$$

where the binomial probability mass function $P_{bin}$ is given by

$$P_{bin}(k, N, p) = \binom{N}{k} p^k (1-p)^{N-k}. \tag{S8}$$

For interval widths $w_A, w_P \ll 1$ we can neglect edge effects, and the resulting pollinator degree distribution will be approximately binomial, i.e.

$$p_A(k) \approx P_{bin}(k, N, p_{int}(w_A + w_P)/(1 - w_P)). \tag{S9}$$

This result is exact when considering only (the individual robustness of) pollinators in the bulk.

#### A. Calculating Temporal Robustness ($a = 1$)

For $a = 1$, the event that a pollinator survives is equivalent to the event that the intervals of the plants it interacts with completely cover its own interval. In order to calculate this we first consider the plant intervals as the nodes of a (random) interval graph, and find the probability that $k$ such nodes, chosen at random, are connected and cover the pollinator interval. The paper [1] considers a similar scenario, where nodes are placed uniformly in $[0, 1]$, and share an edge if they are within a distance $d$ of one another. The probability that, given that 2 nodes are placed at $x$ and $x + y$, a remaining $k - 2$ nodes drawn uniformly from $[0, 1]$ fall inside and cover the interval of width $y$ (so that there is a path between the leftmost and rightmost nodes) is given in lemma 1 as

$$P(k,y,d) = \sum_{j=0}^{\min(k-1,\lfloor \frac{y}{d} \rfloor)} \binom{k-1}{j} (-1)^j (y-jd)^{k-2}. \tag{S10}$$

For a pollinator with active period $[x, x+w_A]$ we can imagine placing two "dummy" plant nodes (plant intervals with midpoints) at $x - w_P/2$ and $x + w_A + w_P/2$, so that the three intervals are placed end to end and there is an "interaction interval" of width $w_A + w_P$ between the two plant nodes. Any further plants will then overlap with the pollinator if and only if their midpoints fall inside the interaction interval. By then rescaling $w_A + w_P \to 1$ so that all further nodes are guaranteed to fall within the interaction interval, the probability that the pollinator interval is covered given it has degree $k$ is given by

$$P(k+2, 1, \frac{w_P}{w_A+w_P}) = \sum_{j=0}^{\min(k,\lfloor \frac{w_A}{w_P} \rfloor)+1} \binom{k+1}{j} (-1)^j (1 - \frac{jw_P}{w_A+w_P})^k. \tag{S11}$$

In this case, the probability that a pollinator is not extinct given that there are $n$ pollinators remaining can therefore be written (using equation S9 and ignoring edge effects)

$$\begin{aligned} p_{s,n}(w_A, w_P) &= \sum_{k=0}^{n} \Pr(\text{survives}|\text{degree} = k) \Pr(\text{degree} = k) \\ &\approx \sum_{k=0}^{n} P(k+2, 1, \frac{w_A}{w_A+w_P}) P_{bin}(k,n,p). \end{aligned} \tag{S12}$$

Here we have also ignored any effects that the edges will have on the covering probability for a given degree $k$.

Using the above equation, the expected robustness be written as

$$\langle R \rangle_{E,S} \approx \frac{1}{N+1} \sum_{n=0}^{N} p_{s,n}, \tag{S13}$$

where $\langle \cdot \rangle_{E,S}$ indicates that we are now averaging over extinction sequences and the network ensemble. By splitting up the sum over $k$, this can be rewritten as

$$\begin{aligned} \langle R \rangle_{E,S} \approx \frac{1}{N+1} &\left( \sum_{n=0}^{N} \sum_{k=0}^{\lfloor\theta\rfloor} \sum_{j=0}^{k+1} + \sum_{n=0}^{N} \sum_{k=\lfloor\theta\rfloor+1}^{n} \sum_{j=0}^{\lfloor\theta\rfloor+1} \right) \\ &\binom{k+1}{j} \binom{n}{k} (-1)^j \left(p - jp'_{int} w_P\right)^k (1-p)^{n-k} \end{aligned} \tag{S14}$$

where $\theta = \frac{w_A}{w_P}$ and $p'_{int} = p_{int}/(1 - w_P)$.

We now introduce the forward difference operator $\Delta$, which can be defined as

$$\Delta f(t) = f(t+1) - f(t), \tag{S15}$$

and note the useful relations

$$\Delta^n f(t) = \sum_{j=0}^{n} \binom{n}{j} (-1)^{n+j} f(t+j), \tag{S16}$$

and, for a degree $k$ polynomial $P_k(t)$,

$$\Delta^n P_k(t) = \begin{cases} 0 & n > k \\ k! & n = k \end{cases}. \tag{S17}$$

By letting $g(t) = (p - tp_{int}w_P)^k$, we can rewrite the left hand sum over $j$ in equation S14 to show that it is proportional to

$$\sum_{j=0}^{k+1} \binom{k+1}{j} (-1)^j (p - jp'_{int} w_P)^k = \Delta^{k+1} g(0). \tag{S18}$$

We can then see from equation S17 that, since $g$ is a polynomial of degree $k$, the leftmost set of summations in equation S14 is equal to 0.

In order to evaluate the 2nd set of summations in equation S14, we first use the identity

$$\binom{n}{m} x^m (x+y)^{n-m} = \sum_{k=0}^{n} \binom{k}{m} \binom{n}{k} x^k y^{n-k}, \tag{S19}$$

which can be obtained by taking the $m$th derivate w.r.t. $x$ of the binomial relation $(x+y)^n = \sum_{k=0}^{n} \binom{n}{k} x^k y^{n-k}$, to evaluate the sum over $k$. Using equations S16 and S17 again we can also prove the identity

$$\sum_{k=0}^{n} \binom{n}{k} (x+k)^k (y-k)^{n-k} = \sum_{\ell=0}^{n} \frac{n!}{\ell!} (x+y)^{\ell}, \tag{S20}$$

which can be rearranged to show that

$$\sum_{j=0}^{\theta} \binom{\theta}{j} (-1)^j \left[ (a-jw)^j (b-jw)^{\theta-j} - (a-jw+\delta)^j (b-jw+\delta)^{\theta-j} \right] = 0, \tag{S21}$$

allowing us to cancel terms in S14. Then, by taking the $m$th derivative w.r.t. $r$ of the geometric sum $\sum_{n=0}^{N} r^n = (1-r^{(n+1)})/(1-r)$ we obtain the identity

$$\sum_{n=0}^{N} \binom{n}{m} r^n = \frac{r^m}{(1-r)^{m+1}} - \sum_{k=0}^{m} \binom{N+1}{k} \frac{r^{N+1+m-k}}{(1-r)^{m+1-k}}, \tag{S22}$$

allowing us to evaluate the sum over $n$. Equation S14 can then be rewritten

$$\begin{aligned} \langle R \rangle_{E,S} \approx 1 - \frac{1+\theta}{p'_{int}(N+1)w_P} \sum_{j=0}^{\lfloor \theta \rfloor} \frac{(\theta-j)^j}{(j+1)^{j+2}} (-1)^j \\ \left[ 1 - \sum_{\ell=0}^{j+1} \binom{N+1}{\ell} (p'_{int}(j+1)w_P)^{\ell} (1-p'_{int}(j+1)w_P)^{N+1-\ell} \right. \\ \left. + \frac{(p'_{int}(j+1)w_P)^{j+2}}{p'_{int}(w_A+w_P)} \binom{N+1}{j+1} (1-p'_{int}(j+1)w_P)^{N-j} \right]. \end{aligned} \tag{S23}$$

We now define $\rho = p_{int} N w_P/(1-w_P)$ as the "effective plant density", which is equal to the expected number of plants that will interact with a pollinator at a given point in time. In the limit $N \to \infty$ such that $\rho$ remains constant and $\theta \ll N$, we can rewrite the previous equation as

$$\begin{aligned} \langle R \rangle_{E,S} \approx 1 - \frac{1+\theta}{\rho} \sum_{j=0}^{\lfloor \theta \rfloor} \frac{(\theta-j)^j}{(j+1)^{j+2}} (-1)^j \\ \left[ 1 - \left( \sum_{\ell=0}^{j} \frac{((j+1)\rho)^{\ell}}{\ell!} + \frac{(\theta-j)((j+1)\rho)^{j+1}}{(1+\theta)(j+1)!} \right) e^{-(j+1)\rho} \right]. \end{aligned} \tag{S24}$$

If $\rho$ is large enough that the exponential term vanishes this can be written more simply as

$$\langle R \rangle_{E,S} \approx 1 - \frac{1+\theta}{\rho} \sum_{j=0}^{\lfloor \theta \rfloor} \frac{(\theta-j)^j}{(j+1)^{j+2}} (-1)^j, \tag{S25}$$

which is equivalent to neglecting the 2nd term in equation S22. We also note that, as $\rho \to \infty$ the robustness tends to 1.

#### B. Calculating Time-Aggregated Robustness ($a = 0$)

The time-aggregated robustness can be found by considering all interactions for a given pollinator to be active across it whole active period, and is equivalent to robustness of a non-temporal network with the same degree distribution. Ignoring edge effects, the pollinator degree distribution is given by the binomial distribution (equation S9). This process therefore produces a network with the same degree distribution as that of a bipartite Erdős-Rényi network, in which each possible edge between a plant and pollinator exists independently with probability (or connectance) $C = p_{int}(w_A + w_P)/(1 - w_P) = p$. Therefore the time-aggregated temporal network has an expected robustness given by

$$\langle R_{\text{agg}} \rangle_{E,S} \approx 1 - \frac{1}{(N+1)p}(1 - (1-p)^{N+1}), \tag{S26}$$

which for $p \ll 1 \ll N$ becomes

$$\langle R_{\text{agg}} \rangle_{E,S} \approx 1 - \frac{1}{Np}(1 - e^{-Np}). \tag{S27}$$

Equation S26 can also be recovered by setting $w_A = 0$ in S23, illustrating that the temporal robustness and the time-aggregated robustness coincide as $\theta \to 0$.

#### C. Calculating Temporal Robustness ($0 < a < 1$)

For intermediate values of $a$ an analytical description has been found for $a\theta < 1$, i.e. where the plant interval $w_P$ is larger than the minimum pollinator breeding period $aw_A$. For $a > 1/2$ the event that the pollinator survives is equivalent to the event that the region of width $(2a - 1)w_A$ directly in the centre of the pollinator is covered, and the robustness becomes that of a uniform interval network with MRT $a' = 1$ and pollinator width $w'_A = (2a - 1)w_A$. This is given by

$$\langle R \rangle_{E,S} \approx 1 - \frac{1 + (2a-1)\theta}{\rho}. \tag{S28}$$

For $a \leq 1/2$ the event a pollinator survives is equivalent to the event that at least one plant interacts with it, with at least $aw_A$ of overlap. The robustness is then given by

$$\langle R \rangle_{E,S} \approx 1 - \frac{1}{\rho(1 + (1-2a)\theta)}. \tag{S29}$$

When $a = 1/2$, these two expressions coincide at $\langle R \rangle \approx 1 - 1/\rho$ and become constant in $\theta$.

### S3. ADDITIONAL DATASETS

In addition to the Donana dataset discussed in the main text, we have access to two others, which amount to 6 temporal networks in total. The first can be found through the studies [2] and [3], in which an arctic plant-pollinator system in northeastern greenland was studied over two years (1996-1997). We note here that the species interactions are inferred through the adjacency matrix presented in [2] (which we assume constant for both years), whereas phenologies for each year are provided in [3]. The other 4 networks are inferred from EuPPollNet [4]. Although this dataset does not provide species phenologies explicitly, the 4 networks corresponding to the study ID "14_Dupont" (gathered in Danish Heathland in 2004-2005, see e.g. [5]) are sampled over many (35-59) days, and so we are able estimate the phenology of each species by taking its first (last) recorded interaction as being the start (end) of its active period. These 6 networks are summarised in table S1.

Here we reproduce results from the main text for these 6 additional networks. Figure S1 is equivalent to panels C and E of Figure 2 in the main text, Figure S2 is equivalent to panel D of Figure 2, and Figure S3 is equivalent to Figure 4. It is worth noting that, in contrast to the Doñana dataset considered in the main text, many pollinators here have an individual robustness of $r = 0$ (for $a = 1$). This indicates that they have gaps in their floral resources (even before node removal) and are thus always extinct in our model, suggesting either incomplete sampling, and/or that species have an (effective) MRT $a < 1$.

| Network Name | No. Plants $N_P$ | No. Pollinators $N_A$ | Sampling Duration | Sampling Days |
|---|---|---|---|---|
| Greenland_1996 | 31 | 61 | 47 | 25 |
| Greenland_1997 | 31 | 64 | 69 | 25 |
| Horbylunde_2005 | 12 | 103 | 139 | 35 |
| Skov_Olesen_2004 | 21 | 142 | 176 | 59 |
| Isen_Bjerg_2004 | 17 | 129 | 190 | 46 |
| Isen_Bjerg_2005 | 14 | 142 | 164 | 39 |

TABLE S1: Summary data of additional plant-pollinators networks

### S4. INFERENCE OF MODEL PARAMETERS FROM DATA

In section IV of the main text we introduce the linear interaction model, in which plant active period durations are randomly drawn from a beta distribution. The beta distribution has a probability density function (PDF) is given by

$$\rho(x) = \frac{x^{\alpha-1}(1-x)^{\beta-1}}{B(\alpha,\beta)} \tag{S30}$$

for $0 \leq x \leq 1$, where $a, \beta > 0$, and $B(\alpha,\beta) = \int_0^1 x^{\alpha-1}(1-x)^{\beta-1}dx$ is the beta function. This PDF is numerically fit, using the `fitdist` function of MATLAB's Statistics and Machine Learning Toolbox (Version 24.1, R2024a), to the plant active periods gathered from data, and is illustrated here for "Skov_Olesen_2004" in figure S4. These inferred beta distributions are then used to randomly generate plant intervals for the linear interaction model.

The linear interaction model also requires a constant of proportionality between the amount of overlap a given plant-pollinator pair has, and their probability of interacting. In order to infer this for a given network, we take each plant-pollinator pair, and plot its interaction probability (i.e., 1 or 0 if they do or don't interact respectively). Using this we perform a linear regression (using MATLAB's `mldivide` function), and take the gradient as the constant of proportionality. This is illustrated in Figure S5 for the "Skov_Olesen_2004" network. Note that, when applied to the linear interaction model, any overlaps that produce an interaction probability $> 1$ were assigned interaction probability 1 (i.e. guaranteed to interact).


[1] E. Godehardt and J. Jaworski, On the Connectivity of a Random Interval Graph, Random Structures and Algorithms **9**, 137 (1996).

[2] J. M. Olesen, J. Bascompte, H. Elberling, and P. Jordano, Temporal Dynamics in a Pollination Network, Ecology **89**, 1573 (2008).

[3] C. Pradal, J. M. Olesen, and C. Wiuf, Temporal development and collapse of an Arctic plant-pollinator network, BMC Ecology 2009 9:1 **9**, 24 (2009).

[4] J. B. Lanuza, T. M. Knight, N. Montes-Perez, W. Glenny, P. Acuña, M. Albrecht, M. Artamendi, I. Badenhausser, J. M. Bennett, P. Biella, R. Bommarco, A. Cappellari, S. Castro, Y. Clough, P. Colom, J. Costa, N. Cyrille, N. de Manincor, P. Dominguez-Lapido, C. Dominik, Y. L. Dupont, R. Feldmann, E. Felten, V. Ferrero, W. Fiordaliso, A. Fisogni, Fitzpatrick, M. Galloni, H. Gaspar, E. Gazzea, I. Goia, C. Gómez-Martínez, M. A. González-Estévez, J. P. González-Varo, I. Grass, J. Hadrava, N. Hautekèete, V. Hederström, R. Heleno, S. Hervias-Parejo, J. M. Heuschele, B. Hoiss, A. Holzschuh, S. Hopfenmüller, J. M. Iriondo, B. Jauker, F. Jauker, J. Jersáková, K. Kallnik, R. Karise, D. Kleijn, S. Klotz, T. Krausl, E. Kühn, C. Lara-Romero, M. Larkin, E. Laurent, A. Lázaro, F. Librán-Embid, Y. Liu, S. Lopes, F. López-Núñez, J. Loureiro, A. Magrach, M. Mänd, L. Marini, R. B. Mas, F. Massol, C. Maurer, D. Michez, F. P. Molina, J. Morente-López, S. Mullen, G. Nakas, L. Neuenkamp, A. Nowak, C. J. O'Connor, A. O'Rourke, E. Öckinger, J. M. Olesen, H. Opedal, T. Petanidou, Y. Piquot, S. G. Potts, E. F. Power, W. Proesmans, D. Rakosy, S. Reverté, S. P. Roberts, M. Rundlöf, L. Russo, B. Schatz, J. Scheper, O. Schweiger, P. E. Serra, C. Siopa, H. G. Smith, D. Stanley, V. Ştefan, I. Steffan-Dewenter, J. C. Stout, L. Sutter, E. M. Švara, S. Świerszcz, A. Thompson, A. Traveset, A. Trefflich, R. Tropek, T. Tscharntke, A. J. Vanbergen, M. Vilà, A. Vujić, C. White, J. B. Wickens, V. B. Wickens, M. Winsa, L. Zoller, and I. Bartomeus, EuPPollNet: A European Database of Plant-Pollinator Networks, Global Ecology and Biogeography **34**, e70000 (2025).

[5] Y. L. Dupont and J. M. Olesen, Ecological modules and roles of species in heathland plant–insect flower visitor networks, Journal of Animal Ecology **78**, 346 (2009).

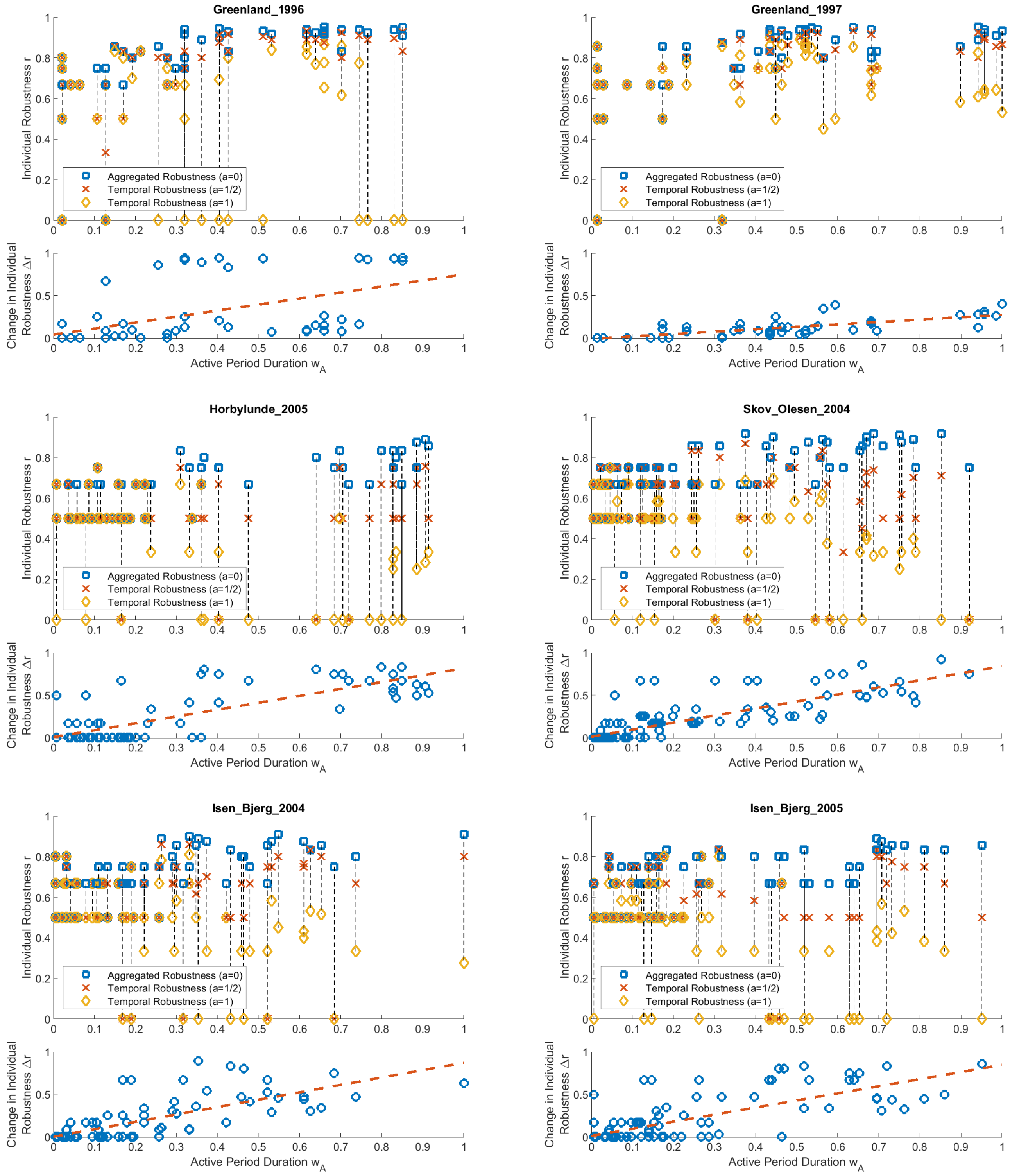


FIG. S1: Equivalent to figure 2 (Panels C and E) of the main text for the 6 networks described in table S1.

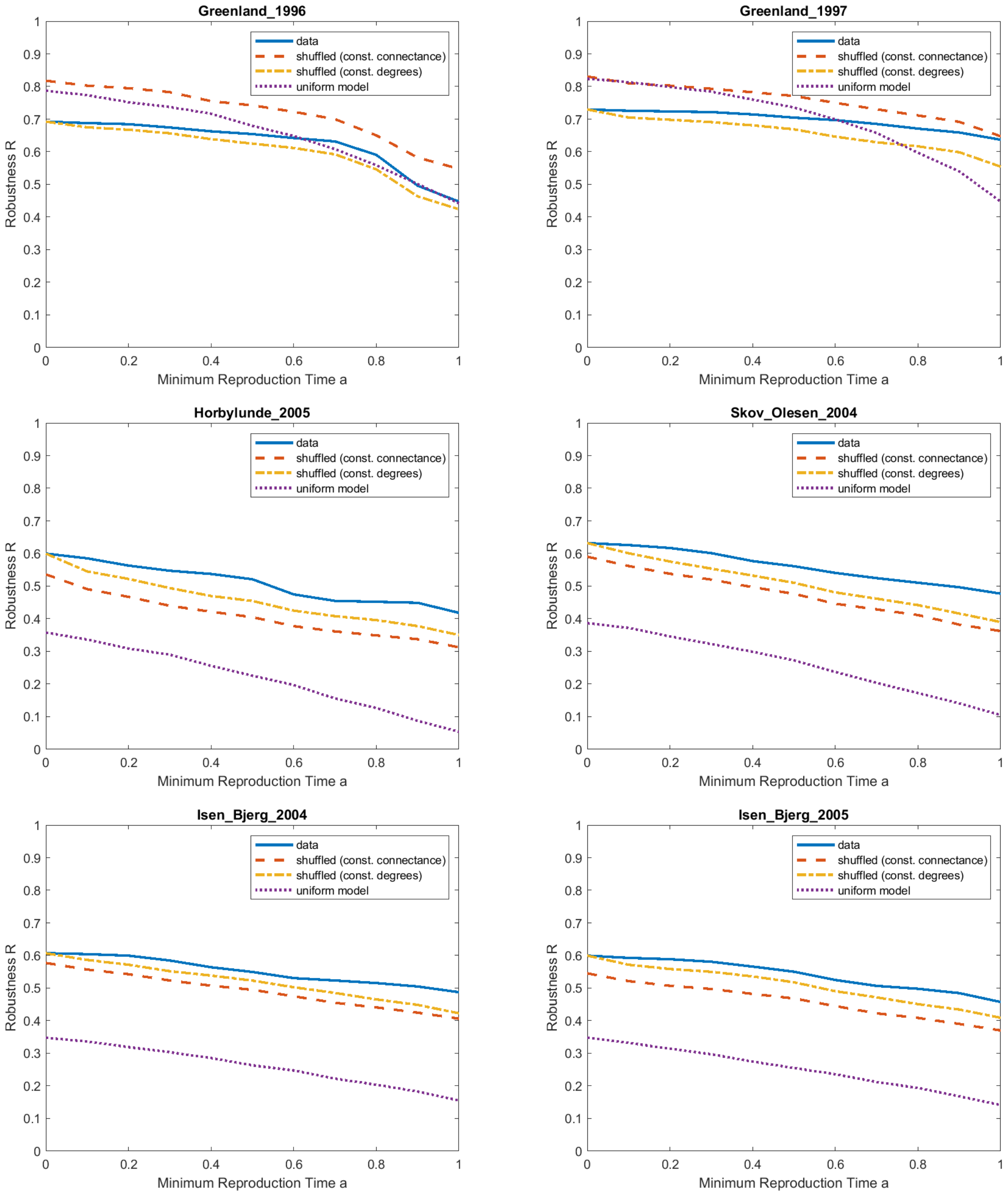


FIG. S2: Equivalent to panel D of Figure 2 of the main text for the 6 networks described in table S1.

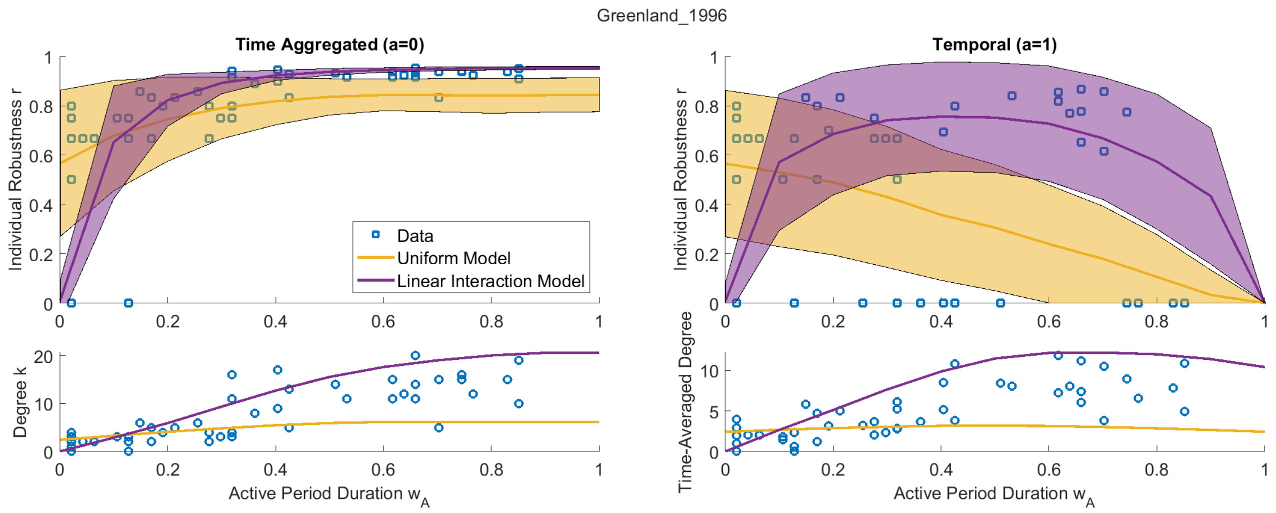
Greenland_1996
Time Aggregated (a=0)
Temporal (a=1)
Individual Robustness r
Data
Uniform Model
Linear Interaction Model
Degree k
Time-Averaged Degree
Active Period Duration $w_A$

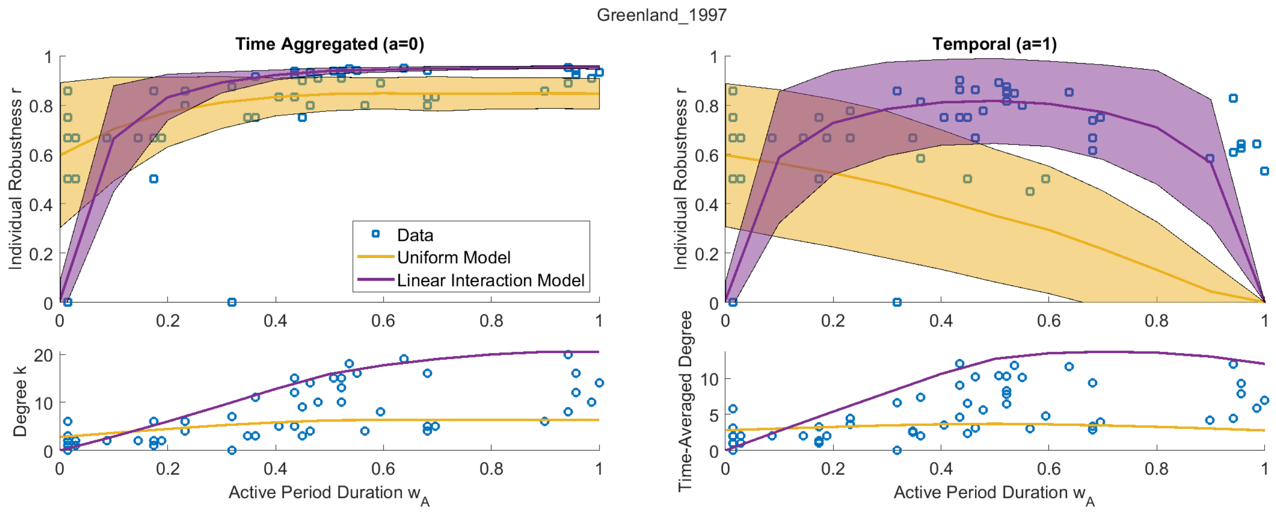
Greenland_1997
Time Aggregated (a=0)
Temporal (a=1)
Individual Robustness r
Data
Uniform Model
Linear Interaction Model
Degree k
Time-Averaged Degree
Active Period Duration $w_A$

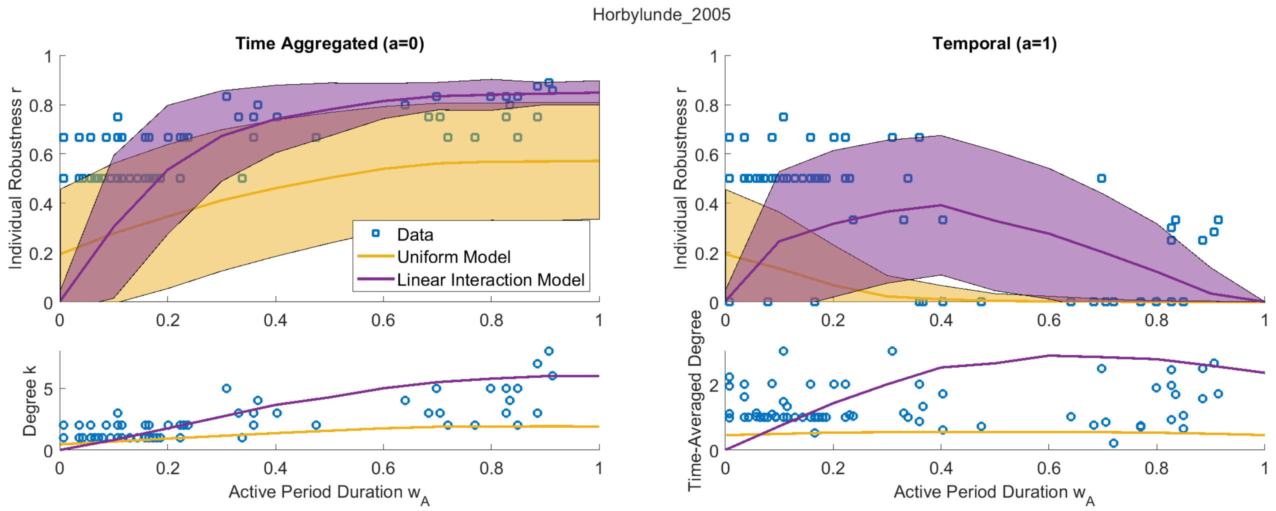
Horbylunde_2005
Time Aggregated (a=0)
Temporal (a=1)
Individual Robustness r
Data
Uniform Model
Linear Interaction Model
Degree k
Time-Averaged Degree
Active Period Duration $w_A$

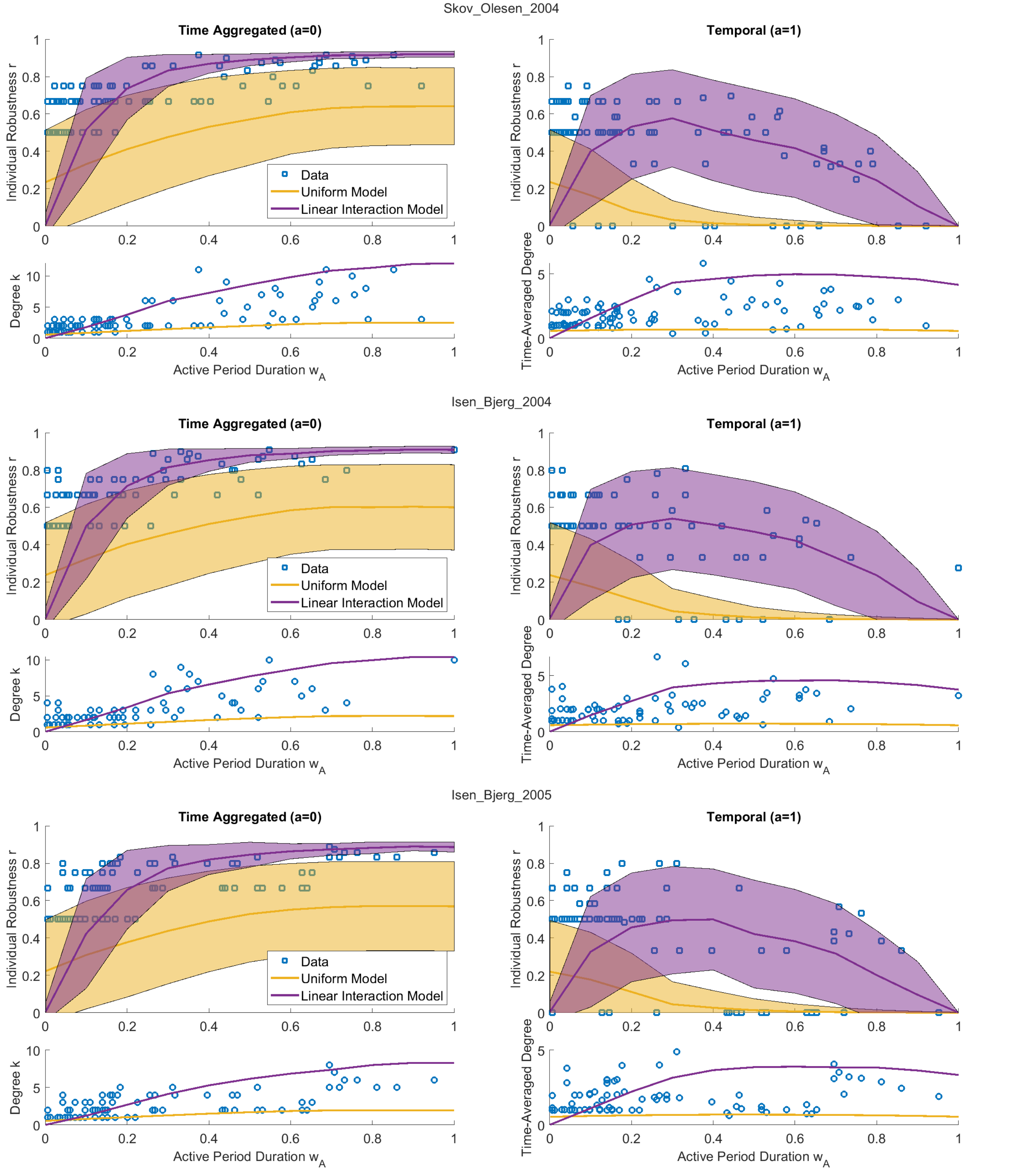


FIG. S3: Equivalent to figure 4 of the main text for the 6 networks described in table S1.

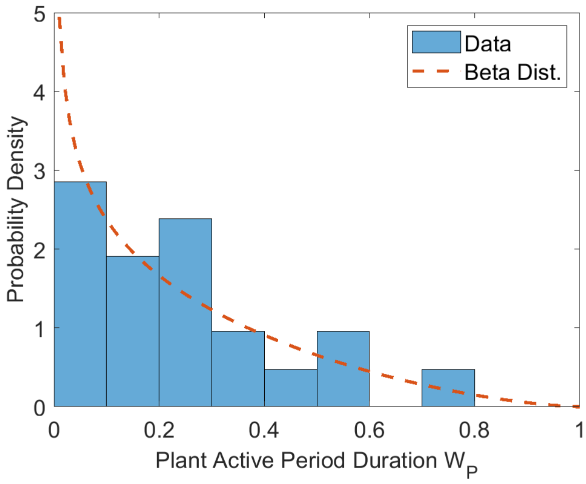


FIG. S4: Example beta distribution fit to model the distribution of plant active period durations in the "Skov_Olesen_2004" network.

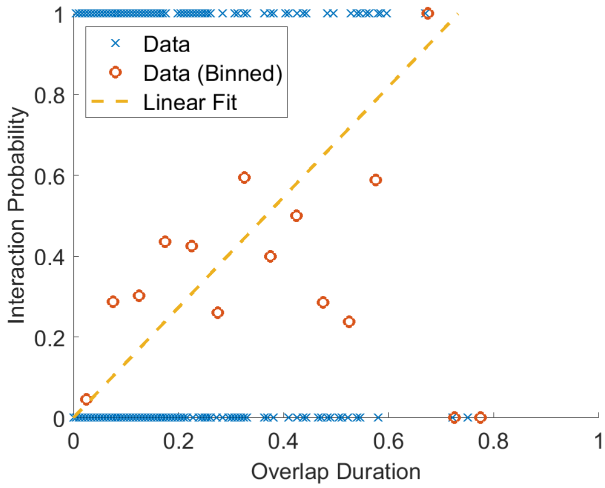


FIG. S5: Interaction probability against overlap duration for the "Skov_Olesen_2004" network. Binned data is the average interaction probability for all plant-pollinator pairs in the network with an overlap within the given bin. Each data point for the binned data is at the midpoint of a bin of width 0.05. Note that the linear fit is fitted to the unbinned data (blue crosses).
